# A spatio-temporaral atlas of the growing brain for fMRI studies

**DOI:** 10.1101/2024.04.15.589560

**Authors:** Maria Murgasova, Valentina Doria, Latha Srinivasan, Paul Aljabar, David Edwards, Daniel Rueckert

## Abstract

Spatio-temporal atlases of the growing brain can help when building models of brain development, not only for anatomical structure, but also for the changing patterns of activation within the brain. In this paper, we build such an age-dependent atlas using registrations among a large number of anatomical datasets, acquired from neonatal subjects of different ages, and followed by kernel-based regression. fMRI activation data are then registered to this atlas to build a model of the development of brain networks, such as the default and motor networks, at this early age.

## 1 Introduction

During the last trimester of pregnancy the brain undergoes a period of rapid growth, for example in the development of cortical folding. Building models of average growth can help to answer many questions about brain development during this phase of growth. A challenge is presented by the data available being scattered between different subjects and different ages. In this paper we describe methodology designed to overcome this challenge and to build such models of growth and development.

Average brain templates using rigid and affine registration [1, 2] have been used for spatial normalization of adult brain MRI. A popular method of building templates using non-rigid registration is to register all the images to a reference image and average anatomy is found by using the inverse of average transformation [3]. An unbiased atlas can be built using group-wise registration using a small deformation model [4] and similarly in a large deformation setting [5]. This approach was extended to building an age-dependent templates using kernel-based regression [6]. The pairwise approach in [3] was also adapted to registration that use free-form deformations (FFDs) [7] and extended to build age-dependent atlases [8].

In this work, we choose the pairwise methodology based on FFD registration as it scales well, allowing additional data to be easily incorporated into the atlas when building models of brain development such as spontaneous brain activation maps which can be extracted from resting state fMRI data [9]. Even though B-spline transformations are not guaranteed to be invertible and optimal in large deformation setting, we assume small deformation, as the resolution of fMRI data is rather low and large deformation is therefore not necessary. The anatomical template of growing brain is built using the methodology described in [8]. Here we propose a method of registering additional data, typically available in smaller quantities, to the age-dependent atlas created using large number of datasets. The data used here is activation data which allow us to build a model of change in brain networks which is representative of the population.

## 2 Creating non-rigid templates

The average brain templates can be created from a large number of anatomical images *I*_1_, …, *I*_*n*_. The images are typically registered into a single reference space *R* via transformations 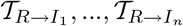. Please note, that for implementational reasons, transformations from the reference to the image are calculated when registering the image to the reference space. If the aligned images, which are defined as 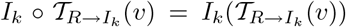, are averaged to produce an atlas, then the atlas tends to be biased towards the reference space and is unlikely to represent the average geometry of the population. Much of this bias can be removed if an average transformation 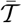 is used to estimate the average space template [3], see Fig. 1.

**Fig. 1.**
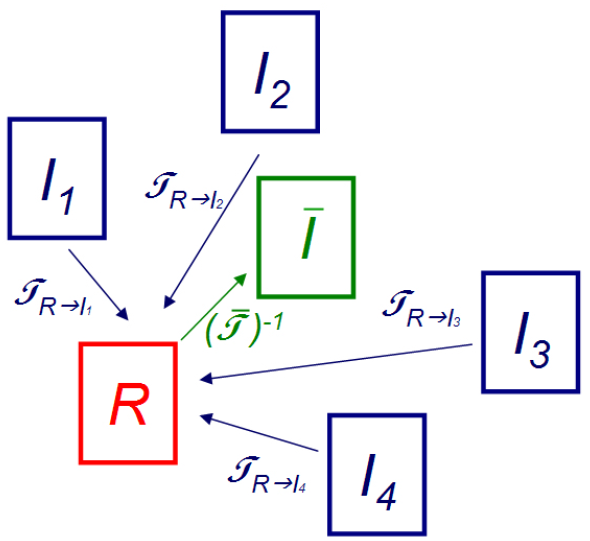
Averaging transformations for atlas creation.

We calculate the non-rigid transformations between brains using FFD registrations that use B-spline basis functions to weight a set of vectors arranged in a regular control point lattice [10].

The general form of the transformations used is

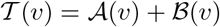

where *v* is a location of a voxel in reference space, *𝒜* is the global affine component and *ℬ* is the non-rigid FFD component. The FFD component is expressed as

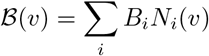

Here *N*_*i*_(*v*) represent 3D tensor-product basis functions and *B*_*i*_ represent B-spline control points.

The neonatal brain undergoes significant global shape change in early neonates. We will, however, focus on changes in activation patterns in this period and consequently, it is desirable not to include the global shape change in the model. We therefore pre-process all the images so that they are affinely aligned with a single reference image. In this case, the transformations 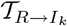 are completely defined by their local FFD component 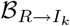 and these are all defined using the same control point lattice in the domain of the reference image. The average transformation 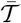 can then be calculated by simple averaging of control points

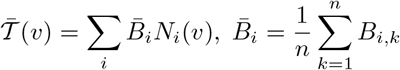

where *B*_*i,k*_, *i* = 1, .., *n* represents the B-spline control points of transformation 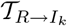.

To define the average space we need to invert the transformation 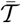. This is again represented by a FFD which is estimated numerically using a Newton method and inverse filtering [11]. Thus the transformation relating the reference to the average atlas space is defined by the transformation 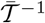 and atlas image *Ī* is calculated as

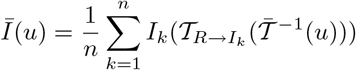

where *u* is a location in the space of the atlas *Ī*.

## 3 Creating spatio-temporal non-rigid atlases

When building a 4D atlas of growing brain, we aim to create a continuous set of 3D volumes or frames dependent on a parameter *t* which, in our case, represents the age of the subjects. This can be achieved by kernel regression, following the work of [6] and [8]. Let *t*_1_, …, *t*_*n*_ be the gestational ages (GA) of the subjects at the time of scan. Then average transformation at the age *t* can be estimated as

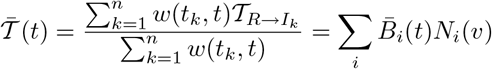

where

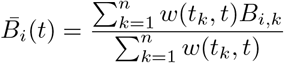

A Gaussian kernel is used to calculate the weights

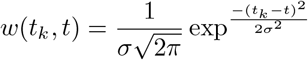

The average template anatomy *Ī*(*t*) is then calculated as

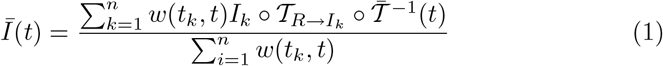

## 4 Propagating data on the spatio-temporal template

Brain networks, such as the default mode or motor networks can be estimated from correlation maps derived from fMRI activation data which are usually acquired together with a corresponding anatomical image. It is possible to use the anatomical images acquired with the fMRI data to create the atlas. In this work, however, we estimate a better representation of population off-line using a large database of anatomical images. The fMRI activation data acquired from a relatively small number of subjects can then be transformed to the space of the 4D atlas by registering their anatomical images to the appropriate frame. We choose this approach over registering the the fMRI data to a common reference or average *Ī*, to minimise the registration error which often increases if the age-difference between the images is large. Let *F*_1_, …, *F*_*m*_ denote the anatomical images with attached correlation maps *C*_1_, …, *C*_*m*_ that represent a brain network. The images *F*_*k*_ can be registered to the template *Ī*(*t*_*k*_) of the corresponding age *t*_*k*_ by non-rigid transformation 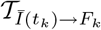. To produce the model 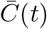 smooth in age *t*, the correlation maps registered to the corresponding frame have to be propagated to the neighbouring frames and averaged for each frame with suitable age-dependent weighting. A transformation from the frame *Ī*(*t*_1_) to the frame *Ī*(*t*_2_) has been calculated while creating the atlas and can be expressed as 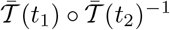. The average correlation map for each age *t* can therefore be calculated as

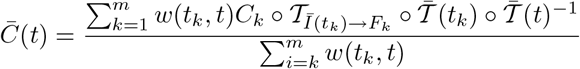

## 5 Implementation and results

To create the 4D anatomical template we used 102 T2-weighted fast-spin echo images acquired on 3T Philips Intera system with MR sequence parameters TR = 1712ms, TE = 160ms, flip angle 90° and voxel sizes 0.86 × 0.86 × 1mm. The age range at the time of scan was 28 to 44 weeks (GA). All subjects were born prematurely. The transformations were estimated in a multi-resolution fashion using control point spacings of 20mm, 10mm and 5mm. We used standard deviation *σ* = 2weeks for calculating the weights. The templates were created with three time-points per week. The resulting template with one frame per week is shown in Fig. 2.

**Fig. 2.**
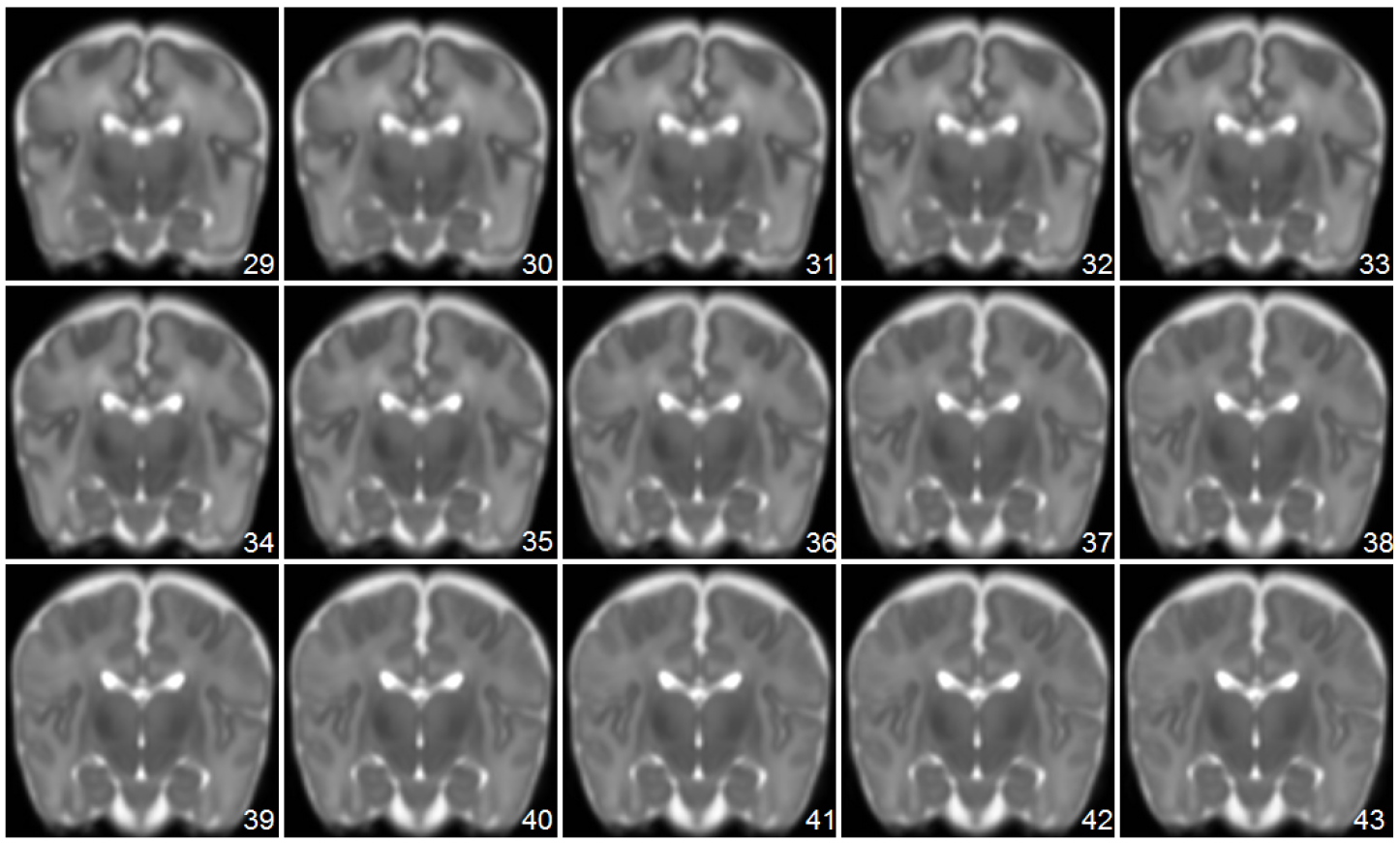
Anatomical template for ages 29-43 weeks GA, obtained by averaging FFDs with a control point spacing 5mm and using a standard deviation of 2 weeks for the Gaussian used in kernel regression.

Spontaneous brain activity can be measured with resting state fMRI. Some brain regions, encompassing neuro-anatomical systems, have been demonstrated to fluctuate in a synchronous manner. Various methods can be used to demonstrate these so-called networks. We focused our attention on two systems: a canonical system, the motor system, and a more controversial system, the so-called default mode network. We used a seed-voxel based functional connectivity analysis: time-series was extracted from a given region (e.g the left motor cortex and the medial prefrontal cortex in our study) and a correlation with the rest of the brain was calculated, producing correlation maps. We processed 39 fMRI datasets acquired from subjects with an age range 29-44 weeks (GA). These consisted of T2 anatomical images and aligned resting-state fMRI data with resolution 2.5 × 2.5 × 3.25mm. The data were processed using FSL [12] to obtain the correlation map estimates representing default mode and motor brain networks. These were then converted to z-scores. The anatomical images were registered to the template as described in Sec. 4 and the average time-dependent model of the developing networks in form of mean z-scores was calculated. Figs. 3 and 4 show the resulting time varying activation images.

**Fig. 3.**
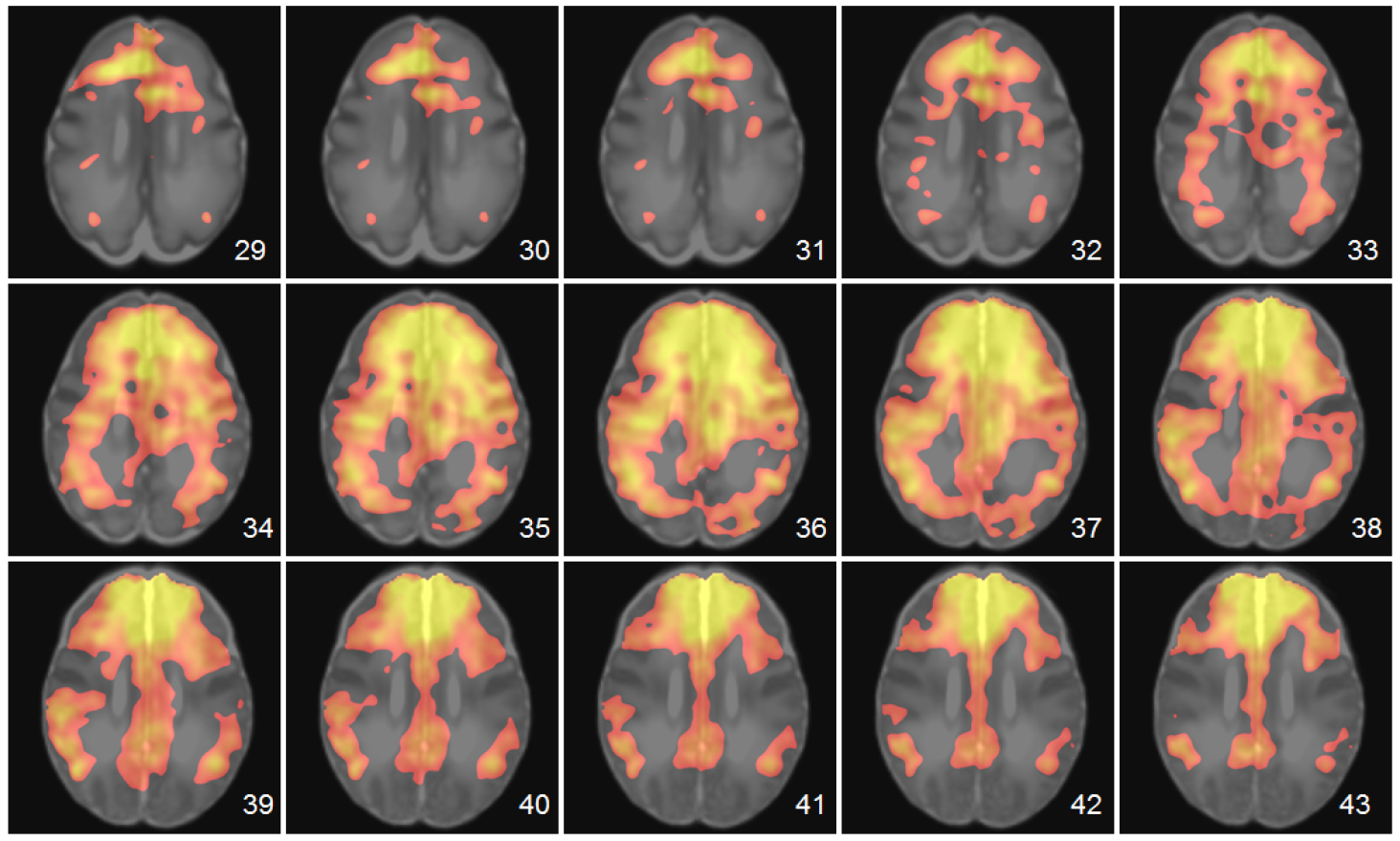
Frames at weekly intervals to illustrate the development of the default mode network for subjects with gestational ages of 29 to 43 weeks.

**Fig. 4.**
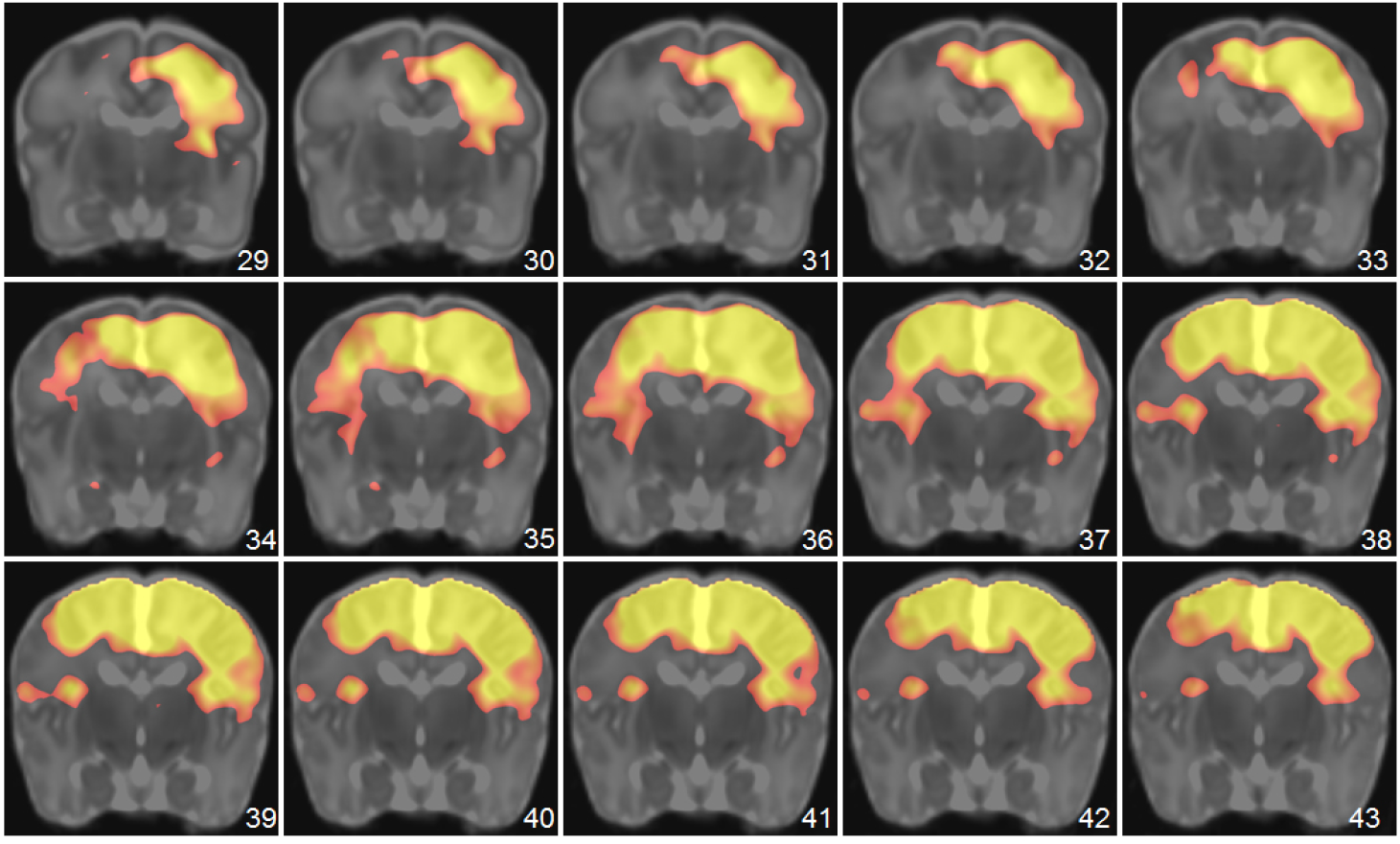
Frames at weekly intervals to illustrate the development of the motor network for subjects with gestational ages of 29 to 43 weeks. The seed was placed in the left motor cortex but the network involves the whole motor system.

## 6 Conclusion

In this work, we have described a method for building a spatio-temporal template of growing brain using FFD based registration and we have shown how such template can be used to estimate spatio-temporal atlases of developing brain networks. We have built the atlases using a small deformation setting, with multi-resolution B-spline FFD registrations with final control point spacing 5mm. We have found that, given the typically low resolution at which fMRI data are acquired, a 5mm control point resolution is sufficient and a finer resolution for atlas creation is unnecessary. The advantage of our approach is that new data can be easily registered and propagated to create new 4D models irrespective of the modality of acquisition as long as an estimate of the anatomy is available.

